# The Neural Consequences of Semantic Composition

**DOI:** 10.1101/2025.08.21.671326

**Authors:** Heather Bruett, Marc N. Coutanche

## Abstract

Humans can create completely new concepts through semantic composition. These ‘conceptual combinations’ can be created by attributing the features of one concept to another (e.g., a *lemon flamingo* might be a yellow flamingo), or drawing on a relationship between concepts (e.g., a *lemon flamingo* might consume lemons). We ask how semantic composition modulates the neural representations of underlying concepts. Combining functional magnetic resonance imaging (fMRI) with multivariate pattern analysis, we interrogate neural patterns for concepts before and after they were subjected to semantic composition. We observe a post-composition shift in neural patterns underlying weakly constrained concepts in the visual system. The composition of strongly constrained combinations draws on a network of semantic regions that include the right inferior frontal gyrus, left angular gyrus, left lateral anterior temporal lobe, and posterior cingulate cortex. Finally, a subset of the semantic network, in left parahippocampal gyrus, distinguishes the manner of composition: relational or attributive. These findings reveal that semantic composition has neural consequences for the composed concepts, and that the manner of composition affects how the brain’s semantic network is deployed.

---

Humans have the ability to create completely novel ideas from familiar concepts. Semantic composition allows us to combine existing concepts in new ways by attributing the features of one concept to another (e.g., *lemon cup* as a yellow cup) or using a novel relationship between concepts (e.g., a cup for holding lemons; Coutanche, et al. 2019; Estes, 2003). The organization of the human conceptual system leads some combinations to be more readily defined than others. Some combinations are less constrained (more ambiguous) because they are: 1) equally likely to be combined attributively or relationally; 2) combined in a way that is not purely attributive or relational. Most studies of semantic composition examine attributive or relational combinations, with stimuli intentionally chosen that are highly likely to be combined in a given way (i.e., strongly constrained). By definition, this cannot reveal the cognitive and neural processes occurring at the time that the type of combination is being selected. Although a weakly constrained combination will ultimately be defined in a way that is attributive or relational, the path for deciding which type of combination to implement is more challenging for these weakly constrained combinations. Here, we examine attributive, relational, *and* weakly constrained combinations with two main questions: 1) How does semantic composition change activity patterns underlying combined concepts? 2) How does semantic composition draw on the semantic network?

Our first goal was to understand how semantic composition influences the brain activity patterns reflecting underlying concepts. How one engages with a concept can influence subsequent processing (Yee & Thompson-Schill, 2016). For instance, directing attention to a concept’s modalities (van Dantzig et al., 2008) or attributes (Bermeitinger et al., 2011) changes subsequent conceptual activation. Similarly, bringing attention to a concept’s color (Yee et al., 2012) or shape (Pecher et al., 1998) induces priming to other concepts with the same feature. Indeed, it has been suggested that concepts should be considered as “changing slightly each time it is retrieved, and that there is no real demarcation between what is activated in a given instance and the concept itself” (p. 1018, Yee & Thompson-Schill, 2016). We hypothesized that because combined concepts involve the manipulation of features or relations from those of its constituent concepts (e.g., *lemon flamingo* might shift the flamingo’s color to yellow), the activity patterns underlying flamingo might in turn be temporarily shifted after composition.

A shift in activity patterns is particularly likely in regions that are influenced by variation in perceptual dimensions, such as visual cortex (e.g., Coutanche & Thompson-Schill, 2019). For instance, naming a concept (in a pre-scan task) subsequently increases adaptation across occipitotemporal cortex (Gotts et al., 2015). A recent investigation identified attentional tuning shifts in occiptotemporal multivariate patterns (via greater discriminability in relevant semantic neighborhoods; Shahdloo et al., 2022), leading to our hypothesis that multivariate patterns would become less similar after attention is focused on particular attributes or relations during semantic composition. We hypothesized that any observed shifts would be particularly strong for weakly constrained combinations, where conceptual attributes or relations undergo greater processing to combine their properties. The simulation account of conceptual combination suggests that simulation mechanisms play a central role in the compositional process (Wu & Barsalou, 2009). Wu and Barsalou (2009) found that people spontaneously generate item properties during conceptual combination that are very similar to those generated during deliberate imagery. The perceptual properties of mental images affect the amount of processing involved (Shepard & Cooper, 1982), such that “just as it takes longer to scan across 30° of visual angle in the visual field than 10°, it also takes longer to scan across 30° of a visual image than 10°” (p. 174, Wu & Barsalou, 2009). Weakly constrained combinations are likely to involve additional simulation processing since the attribute and relational space is explored further to reach a final combination solution.

A second goal was to understand how the semantic network is activated by different forms of conceptual combination: strongly constrained (relational, attributive) and weakly constrained. Strongly constrained relational and attributive combinations are thought to have unique neural underpinnings (Estes, 2003; Wisniewski, 1997), so we hypothesized greater activation for attributive combinations (where a perceptual feature of the modifier noun is shifted to the head noun; e.g., yellow from lemon to flamingo) in regions that bind perceptual properties, such as the anterior temporal lobe (ATL; Boylan, et al. 2017; Coutanche & Thompson-Schill, 2015). Concrete nouns produce a network of bilateral activation featuring both the left and right ATL (unlike abstract nouns, which activate a more left-sided network; Sabsevitz et al., 2005). Additionally, both the left and right ATL are thought to hold modality-invariant representations of semantic concepts (Visser & Lambon Ralph, 2011), suggesting that each may be active during conceptual combination. The angular gyrus (ANG) is another key region of the semantic network (Lambon Ralph et al., 2017). The left ANG is responsive to the semantic richness of words (Ferreira et al., 2015), while we have found that the right ANG provides semantic feedback to perceptual regions (Coutanche & Thompson-Schill, 2019), giving reason to suggest that left and right ANG might show greater activation for binding strongly constrained compounds (which combine readily retrieved semantically rich information). We also hypothesized that the competitive selection of semantic information for weakly constrained combinations would lead to greater activation of the left inferior frontal gyrus (IFG; Grindrod et al., 2008; Solomon & Thompson-Schill, 2017; Thompson-Schill, et al. 1997). We also examined the right IFG, which has been reported as being active alongside the left IFG in studies of semantic unification (Zhu et al., 2012) and is important for targeting attention to conceptual features (Hampshire et al., 2009), which is required when creating and selecting a conceptual combination.

In this study, we measured neural activity using functional magnetic resonance imaging (fMRI) as individuals processed familiar concepts before, during and after semantic composition. We ask how the brain combines known concepts, and identify the neural consequences for recently combined concepts.

## Materials and Methods

### Participants

Data were collected from 25 participants based on a prior study of 24 participants (with an additional one-subject buffer) that was able to detect semantically-driven shifts in multivariate patterns (Coutanche & Thompson-Schill, 2019). All participants were screened prior to participation. They were included in the study if they: 1) were between the ages of 18 and 35, 2) indicated they were a native speaker of American English, 3) had no learning or attention disorder, 4) were right-handed, and 5) had no MR contraindications. Two participants were removed for excessive head motion (maximum between-TR head movement greater than the voxel size; 2mm), leaving us with a final sample of 23 participants (age *M =* 22.6; 14 female, 9 male). All study procedures were approved by the Institutional Review Boards at the University of Pittsburgh and Carnegie Mellon University and participants provided written informed consent prior to participation.

### Experimental Design

A summary of the procedure can be found in Figure 1. Participants began with an anatomical scan. They then completed a series of tasks during functional runs: Pre-combining (‘Pre’), Combining, and Post-combining (‘Post’). Participants began with two Pre runs where they passively viewed the head nouns from the word pairs. Each trial consisted of 4 s of viewing and trials were separated by a jittered ITI, lasting an average of 2 s. Each of the 24 head nouns was repeated three times in each Pre run (72 trials). Next, during three Combining runs, participants were presented with novel conceptual combinations featuring the same head nouns and were asked to imagine a new meaning. Participants were told that the meanings they were imagining had no right or wrong answer and that they should simply mentally define the combination as they saw best. Participants pressed a button to indicate when they had decided on the meaning of the word pair. Each trial consisted of 1 second where the noun pair appeared, followed by 5 s to imagine the definition. Trials were separated by a jittered ITI lasting 2 s on average. Each of the 24 noun pairs was presented once per run. After combining, participants completed two Post runs, again passively viewing the same head nouns from Pre and combining runs. The format was consistent with that of the Pre runs, where each trial lasted 4 s to view the head noun followed by a jittered ITI with an average length of 2 s. Head nouns were repeated three times each run.

**Figure 1.**
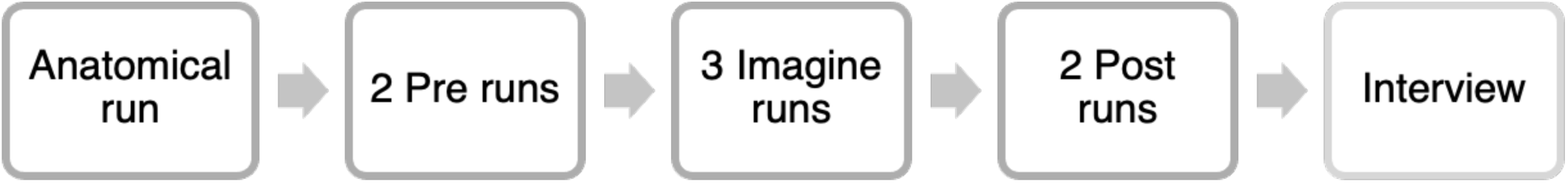
Procedure overview.

### Stimuli Selection

The 24 compounds comprised eight attributive, eight relational, and eight weakly constrained combinations, based on norming in a separate sample of 60 participants. Initial word pairs were selected from past studies: attributional word pairs (*n* = 48) from Estes (2003); relational pairs (*n* = 58) from Estes (2003) and Wisniewski & Love (1998); and weakly constrained pairs (ambiguous; *n* = 82) from Gagné & Shoben (1997) and Kenett & Thompson-Schill (2017). Words were removed for containing compound words (e.g. sleeping pill), having matching taxonomic categories between words, having concreteness levels below a 4 according to the Brysbaert Concreteness Ratings, leaving 34 attributive, 41 relational, and 48 weakly constrained pairs for norming. For norming, we collected data from 60 participants through Prolific (https://prolific.co/). Each participant viewed a random selection of 30 pairs (of the 123 selected earlier). For each pair, participants were asked to provide: their preferred definition for the combination, 1 to 7 Likert scale response for their agreement with two statements (i.e., “I had trouble thinking of a definition for this word combination.”; “I feel that multiple definitions could apply to this word combination equally well.”), and if they were unfamiliar with either word. Participants who indicated that they had a neurological disability, intellectual or learning disability, and/or attentional disorder, or did not meet our language requirements, were removed, giving 55 participants in the final norming sample. Three independent raters categorized the norming participants’ definitions as attributional, relational, neither, impossible to determine, or unsure. Raters were trained on identifying relational and attributional combinations in advance: “attributional” if one dominant attribute or feature from the modifier noun was applied to the head noun, and “relational” if a relationship was formed between the nouns. “Neither” was used to categorize valid definitions where words were combined without attributional or relational techniques (e.g., using the second noun to modify the first; e.g., describing an *elephant ant* as a small elephant). Definitions were categorized as “impossible to determine” if they did not have enough detail for another classification. Final categorizations were determined in a winner-takes-all manner from the 3 independent raters. A total of 24 nouns were selected (8 attributional, 8 relational, 8 weakly constrained). All attributional and relational nouns had at least 75% of definitions categorized as attributional or relational, respectively. Weakly constrained nouns were selected if they were categorized as below 60% attributional and relational. The final stimuli list can be seen in Table 1.

**Table 1.** Stimuli pairs.

| **Attributional** | **Relational** | **Weakly constrained** |
| --- | --- | --- |
| leech boyfriend | book string | attic leg |
| molasses traffic | bowling sweater | cake confetti |
| octopus chair | concrete fountain | copper horse |
| piranha lawyer | floor television | grease fish |
| rocket sprinter | microwave sandwich | olive signals |
| sedative voice | patio cigarette | onion bus |
| strawberry ink | prisoner graffiti | picture soup |
| thunder applause | rodeo magazine | trash house |

### Post-scan

After scanning, participants completed an interview in which they gave definitions of the combinations they had created during the scan, and discussed any strategy used. The session ended with a short survey delivered on Qualtrics, where participants indicated how easy/challenging (1 = very easy, 7 = very challenging) each was to combine, and maintain a single definition. Participants were then given an opportunity to type-out any additional definitions they might have subsequently remembered. Participants then completed a Flanker task, Vividness of Visual Imagery Questionnaire (VVIQ), and semantic test (not analyzed here), before finishing with a brief demographic and language history questionnaire (adapted from Tokowicz, et al. 2004).

### Data Acquisition

Participants were scanned using a Siemens 3-T Prisma magnet and standard radio-frequency coil equipped with a mirror device to allow for fMRI stimuli presentation. Whole-brain imaging was conducted. T1-weighted images were acquired at the start of both sessions (TR = 2.3 s, TE = 1.99, voxel size = 1.0 × 1.0 × 1.0 mm). All five functional runs (TR = 2.0 s, TE = 30) employed voxel sizes of 2.0 × 2.0 × 2.0 mm. A predetermined jitter (average of 2 s) was used in all functional runs, as determined using Optseq 2 (Greve, 2002).

### Data Preprocessing

Imaging data were preprocessed using the Analysis of Functional NeuroImages (AFNI) software package (Cox, 1996). Anatomy were skull stripped and deobliqued using AFNI’s @SSwarper. Functional data were slice time corrected and motion corrected to register them to the third functional volume. Runs were scaled to a 0 – 200 range with a mean of 100 for each voxel and a highpass filter was applied to remove low-frequency trends below .008 Hz from all runs. For univariate analyses, data were smoothed using a kernel with full width with a half maximum (FWHM) of 6 mm. For multivariate analyses, betas were calculated separately for the Pre runs and Post runs, using the Least Squares-Separate (LSS) method based on the onset time of the image appearing on the screen (Mumford et al. 2012).

### Regions of Interest

The ATL ROI was defined with a 10 mm-radius sphere in each hemisphere within MNI space (43 -13 -25; -42 -13 -23; Coutanche & Thompson-Schill, 2015) and then warped to each participant’s native space. The ANG ROI was created in FreeSurfer, as was the IFG, which was formed by combining the pars orbitalis, pars opercularis, triangularis, and the most dorsal region of the IFG (Hirshorn & Thompson-Schill, 2006).

### Statistical Analysis

#### Univariate

Orthogonal contrasts were created to compare strongly constrained combination types (attributive, relational) with weakly constrained, and to compare attributive with relational. Analyses were first conducted in pre-determined ROIs. They were conducted separately in the left and right hemispheres for each ROI: ATL, ANG, and IFG. To determine significance of the contrasts, the average beta coefficient for each contrast was calculated in each subject in each ROI. An independent t-test was used to determine whether the average beta was significantly different from zero. A group-level cluster analysis was also run to determine whether any other regions of the brain showed significant activation differences for our contrasts. To do this, the beta coefficients calculated in native space were warped into standard MNI space. False discovery rate (FDR) correction thresholds were calculated using AFNI’s 3dttest++ with the ClustSim option.

In addition to analyses of activation levels, we also conducted a time-to-peak analysis in the ATL based on Boylan et al. (2017)’s finding of combinatorial differences in the ATL’s timecourse. Analyses were conducted using AFNI’s TENTzero function to calculate the finite impulse response (FIR) for each condition in each voxel (Boylan, et al. 2017) in the right and left hemispheres of the ATL. The TENT function was implemented with a bin-width equal to the 2 s TR and with 10 knots representing 9 intervals ((6 s trial + 12 s HRF)/ 2 s bin-width) modeled for each trial. Given that the FIR models each knot and zeros out the first and final knot, it output 8 beta coefficients for each voxel for each condition. Latency of each subject’s effect peak was determined for each condition, separately for the left and right ROI, as whichever time point had the highest magnitude. Significant time-to-peak differences between attributive and relational conditions were determined by conducting nonparametric Wilcoxon signed rank tests.

#### Multivariate

Betas were calculated separately for each word pre- and post-combination, using LSS. To determine the extent to which activity patterns changed as a result of the combination phase, we conducted a searchlight analysis, in which 8mm-radius searchlights were sequentially examined by correlating their vectors of betas pre- and post-combining. Correlations were Fisher-Z transformed, averaged across words of the same combination type (i.e., weakly constrained, relational or attributive), and subjected to general linear tests comparing strongly constrained combinations to weakly constrained, and attributive to relational. Residual error from the model was used to determine cluster thresholds using AFNI’s 3dClustSim, with bisided selected (tails are tested independently; positive and negative values are not grouped together in the same cluster) and NN=3 (face, edge and cornerwise neighbors for cluster voxels). Significant clusters surpassed a p < 0.005 threshold with cluster-correction bringing the overall alpha to p < 0.05.

## Results

### Behavioral results

Consistent with the stimulus norming, the post-scan interview data indicated that ‘attributional’ combinations were more likely to be combined in an attributional than relational manner (attributional: 83.9%; relational: 10.7%; other: 5.4%), ‘relational’ combinations were more likely to be combined in a relational than attributional manner (relational: 79.5%; attributional: 9.8%; other: 10.7%), and ‘weakly constrained’ combinations were more evenly divided between the two (attributional: 50.0%; relational: 34.8%; other: 15.2%). The post-scan survey confirmed that the weakly constrained combinations (M = 2.72) were significantly more “challenging to define” than the strongly constrained combinations (t_22_ = 2.85, p = 0.009) with no significant difference between relational (M = 1.95) and attributional (M = 2.05) combinations (t_14_ = -0.59, p

= 0.56). Similarly, participants reported it was significantly more challenging to stick to one definition for the weakly constrained combinations (M = 2.16) than the strongly constrained combinations (t_22_ = 2.44, p = 0.02), with no significant difference between relational (M = 1.84) and attributional (M = 1.79) combinations (t_14_ = 0.29, p = 0.77). When asked to list any alternative definitions that came to mind, participants were significantly more likely to list alternatives for weakly constrained (M = 0.42) than strongly constrained combinations (t_22_ = 2.29, p = 0.03), with no significant difference between relational (M = 0.34) and attributional (M = 0.33) combinations (t_14_ = 0.31, p = 0.76).

### Examining neural representational shifts after conceptual combination

Our first question was whether and how semantic composition modulates the neural activity of concepts. We used an exploratory searchlight to identify regions with different levels of neural representational change based on the type of semantic composition: attributional, relational, weakly constrained. One region was identified –left early visual cortex (EVC; MNI center of mass: x = 18; y = 91; z = -4)– where weakly constrained combinations produced greater change in the respective concepts’ activity patterns than did strongly constrained combinations (Figure 2). A multivariate pattern shift was not observed in the ROIs of the conceptual combination task (p > 0.25).

**Figure 2.**
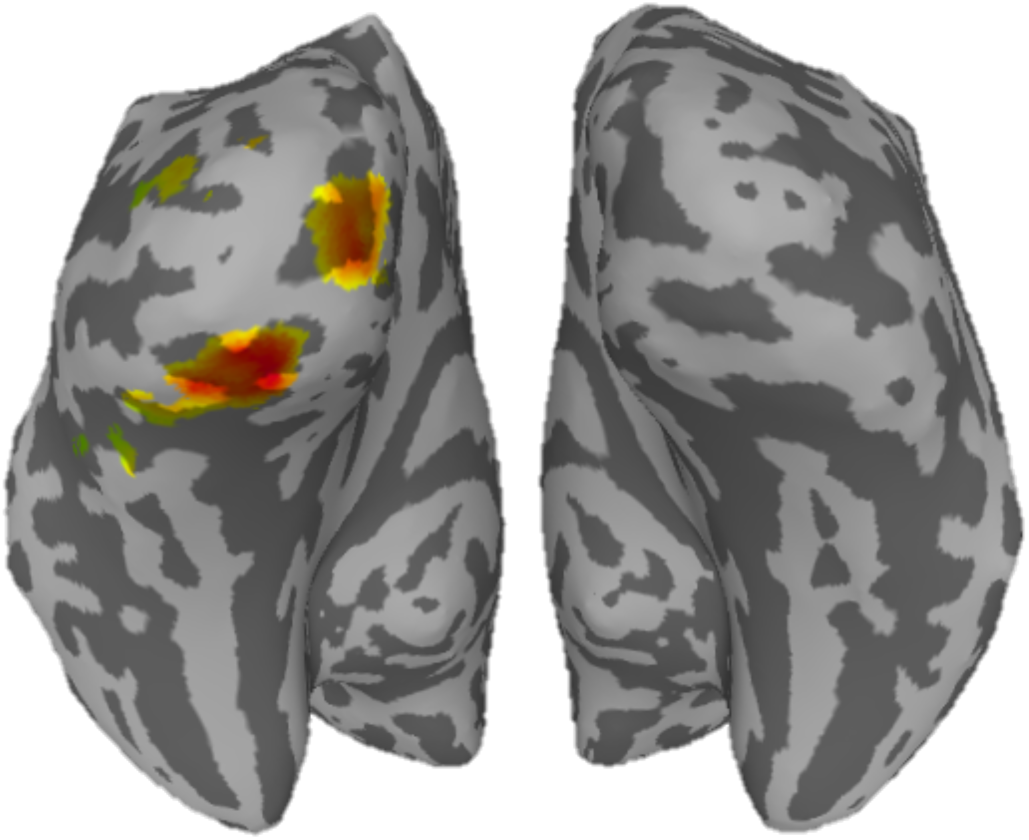
Searchlights with greater shifts in activity patterns after weakly constrained than after strongly constrained combinations. Colors reflect searchlights (8mm radius) with greater shifts for weakly constrained than for strongly constrained word pairs, measured as the correlation between beta coefficients underlying the same concept before, and after, conceptual combination. The correlations are assigned to each searchlight’s central voxel before being Fisher Z-transformed and subjected to a general linear test across the group.

### Antecedents of shift

In order to understand the semantic composition process, we asked how different types of combinations differentially draw on the brain’s systems during the combinatorial process. Comparing strongly constrained and weakly constrained combinations allowed us to identify the neural activity responsible for an easy transference of attributes or relationship versus selecting attributes or relationships through competition. Of the three examined ROIs (ANG, ATL, IFG), the right IFG (*t*(22) = 2.40, *p* = .03) and left ANG (*t*(22) = 2.05, *p* = .05) showed greater activation for strongly constrained relative to weakly constrained combinations (Figure 3, left). For the more specific contrast of attributive and relational combinations, the left IFG was more active at a trending level for attributive than for relational combinations (*t*(22) = 1.90, *p* = .07; Figure 3, right).

**Figure 3.**
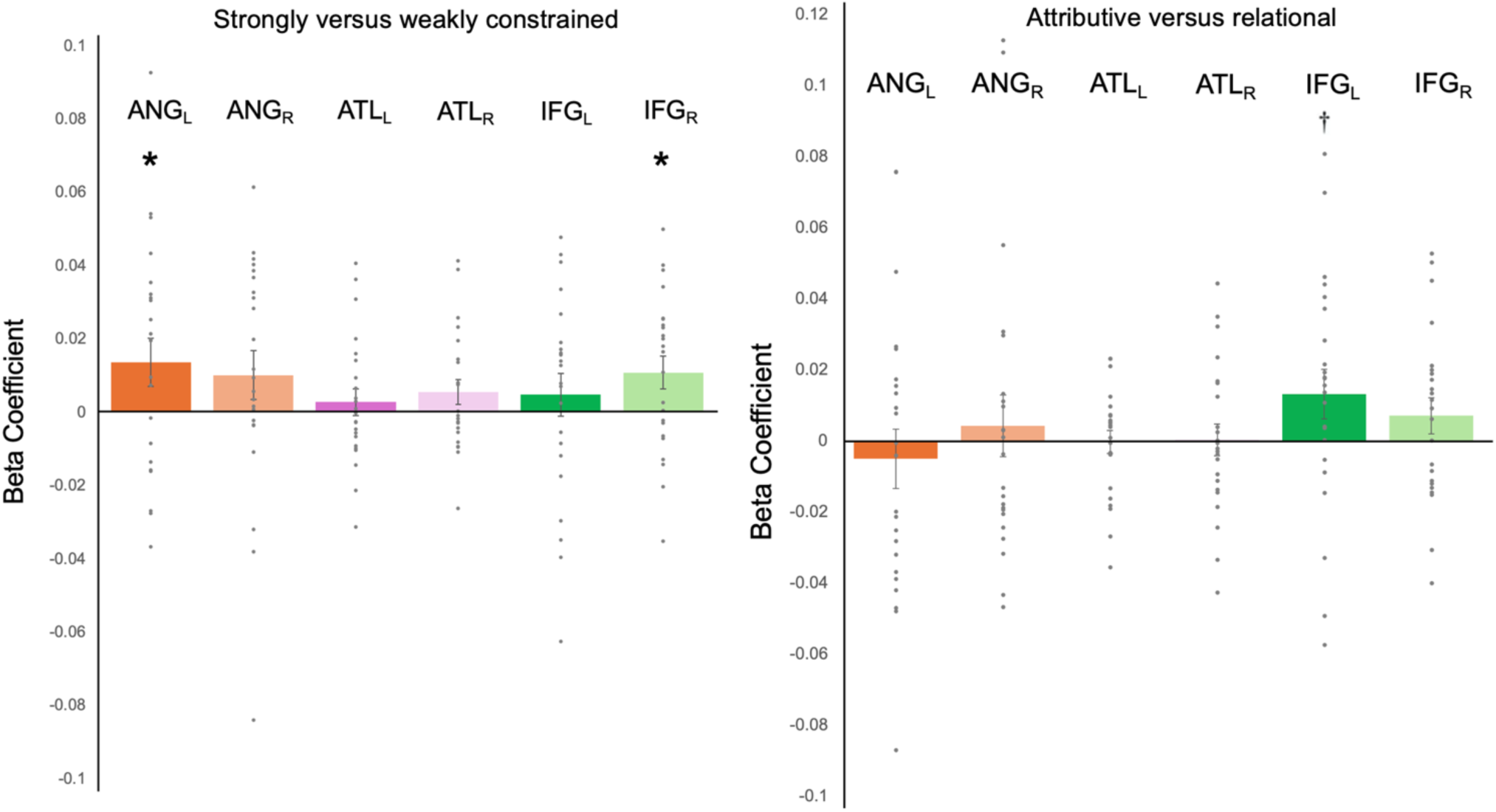
ROI activation. Left: strongly constrained (positive) vs. weakly constrained. Right: attributive (positive) vs. relational. Values reflect average activation level for each ROI during the conceptual combination task, grouped by type of combination. An asterisk indicates statistical significance in an independent t-test comparing the values to zero. A cross indicates a trend. ANG = Angular Gyrus; ATL = Anterior Temporal Lobe; IFG = Inferior Frontal Gyrus; L = left; R = right.

In prior work, we found that conceptually driven activity patterns in occipital regions were accompanied by modulation of left ATL (Coutanche & Thompson-Schill, 2015) and right ANG (Coutanche & Thompson-Schill, 2019). We next asked if these regions’ activation during the conceptual combination process predicts the magnitude of the multivariate pattern shift observed in the occipital cortex clusters. Specifically, we correlated the participants’ mean pattern shift (weakly – strongly constrained; shown in Figure 2) with the corresponding ‘weakly > strongly constrained’ beta values from the intermediary conceptual combination task (shown in Figure 3). This showed that participants with greater occipital pattern shifts after weakly constrained combinations had significantly greater ‘weakly > strongly constrained’ activation during the combination task in the left ATL (r = 0.526, p = 0.01), and significantly lower ‘weakly > strongly constrained’ activation (i.e., strongly > weakly) in right ANG (r = -0.445, p = 0.03).

### Time-to-Peak Analysis

Based on prior work suggesting latency differences in ATL’s timecourse (Boylan et al., 2017), we examined the time-to-peak in the right and left ATL for relational and attributive combination strategies. Boylan et al. (2017) did not find a combinatorial effect in ATL BOLD activation magnitude but did observe differences in the *timecourse* of ATL BOLD response. Specifically, they observed an earlier ATL peak induced by attributive, compared to relational, interpretations. For this reason, we examined the response curves for left and right ATL for the different types of combination. Neither region showed significant peak latency differences (*p*s > .322; Figure 4).

**Figure 4.**
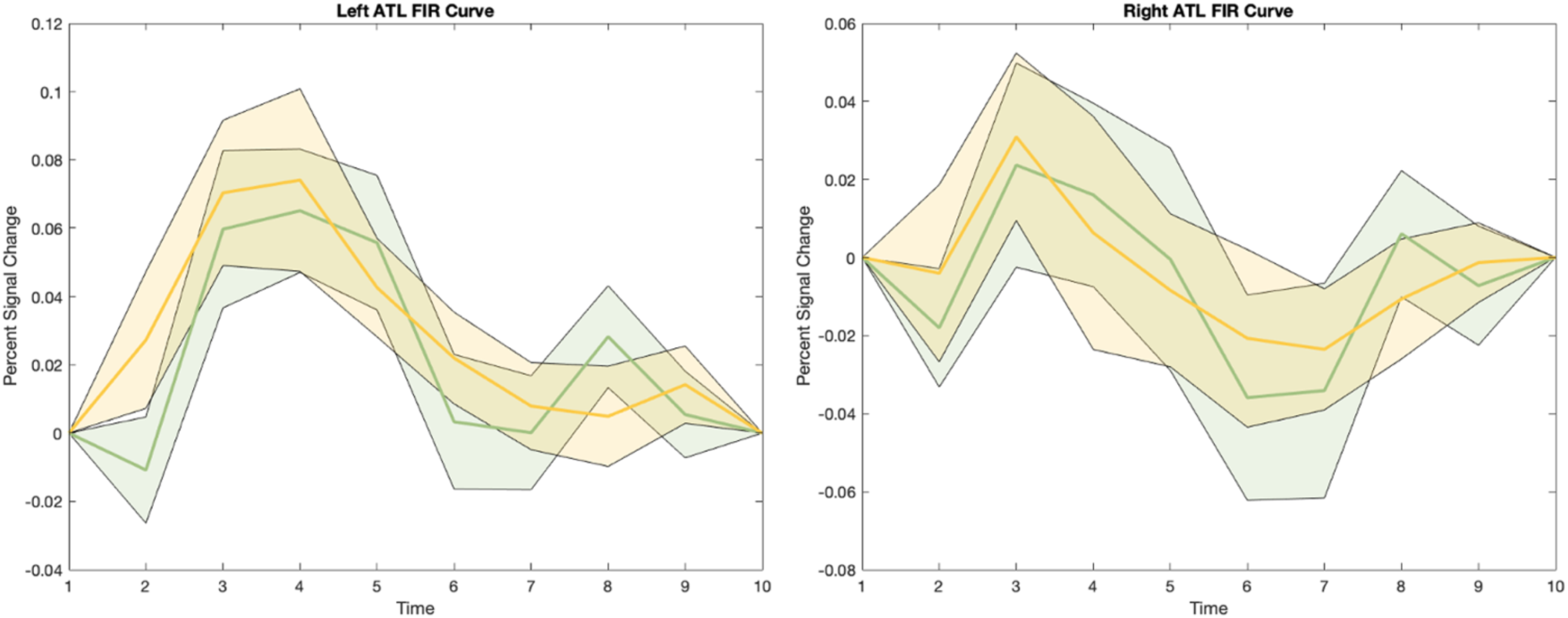
FIR curves of left and right ATL. Signal response curves reflecting percent signal change over time for attributive (green) and relational (yellow) conceptual combinations during the conceptual combination task in left (left panel) and right (right panel) ATL. Error bars indicate +/-1 SE.

### Wholebrain analysis

In addition to the planned ROIs, we ran a wholebrain cluster analysis of the same contrasts. This revealed greater activation in the left ANG, posterior cingulate cortex, and a lateral sub-region of the left ATL (more lateral and ventral than our left ATL ROI) for strongly constrained, relative to weakly constrained, combinations. One cluster in the parahippocampal gyrus was more active for relational than for attributional semantic composition (Table 2; Figure 5).

**Table 2.**
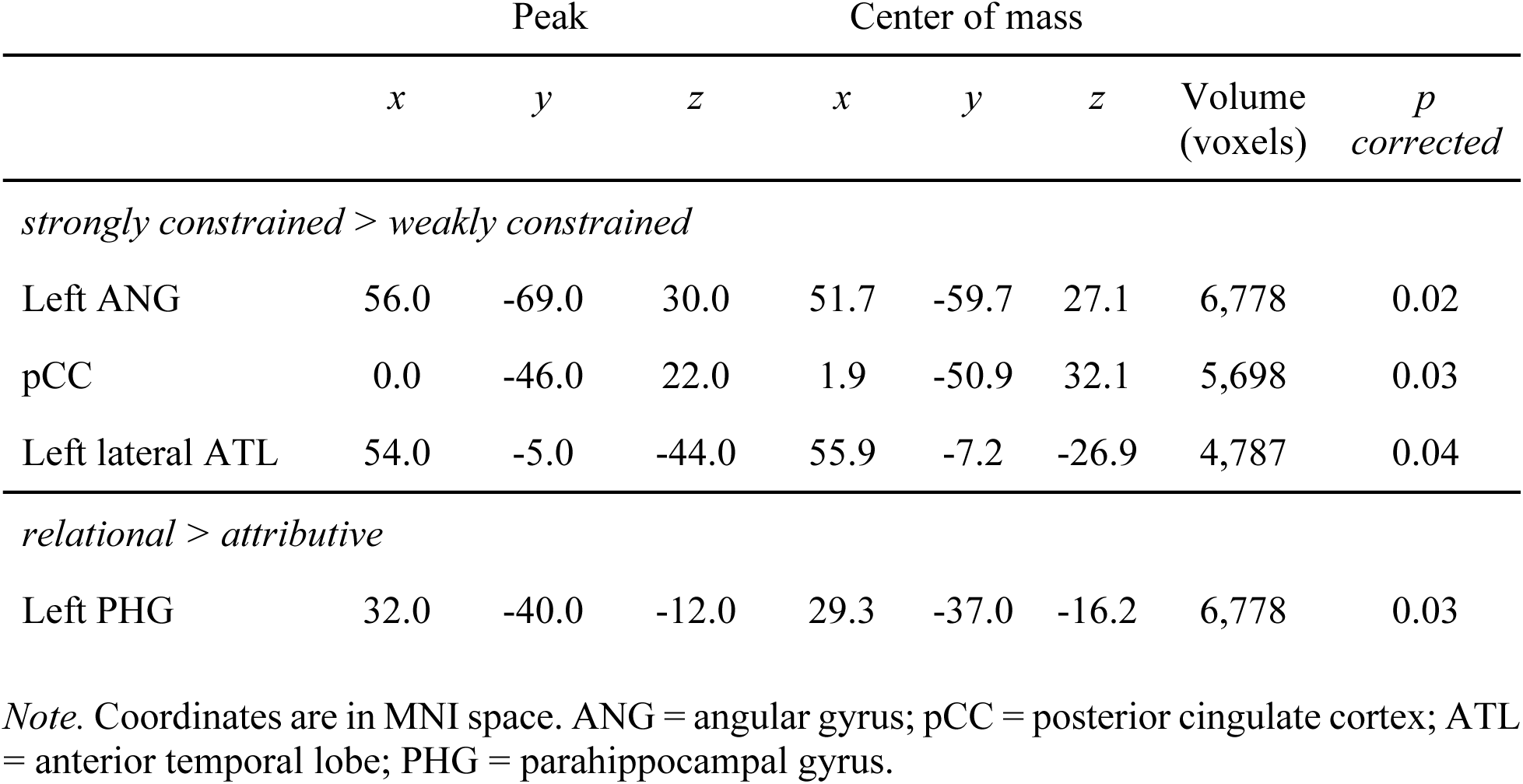
Significant clusters from wholebrain cluster analysis.

**Figure 5.**
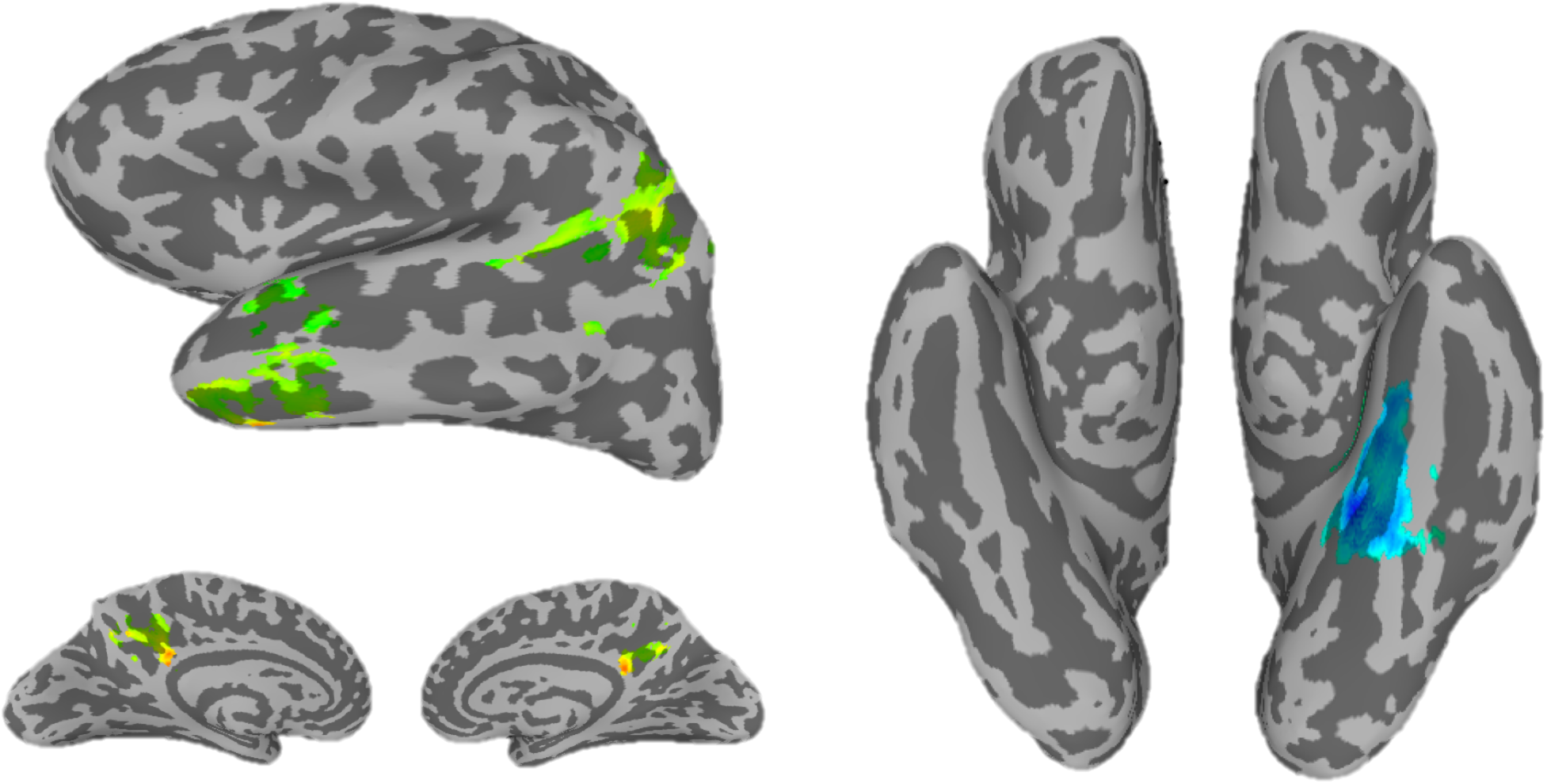
Results of univariate analyses. Left: clusters where strongly constrained combinations showed significantly greater activation than weakly constrained combinations during the conceptual combination process. Right: clusters where relational combinations showed significantly greater activation than attributive combinations during the conceptual combination process. Significance reflects the output of a wholebrain cluster analysis of a group-level t-test against zero with FDR correction.

## Discussion

This study examined the neural underpinnings of semantic composition, and the neural representational changes that semantic composition induce, with a focus on how this differs based on the type of combination.

### Shifting neural representations

This study found a greater shift in neural activity patterns after concepts were combined in weakly constrained, compared to strongly constrained, forms. This shift was focused in areas of visual cortex. In a recent study of selecting targets in preparation for visual search, Witkowski and Geng (2023) identified visual cortex, dorsolateral prefrontal cortex and inferior frontal junction as forming an interconnected network representing the target identity. Analogously, participants in our study were selecting a conceptual identity during conceptual combination. Another recent study identified that occipitotemporal multivariate patterns undergo task-dependent attentional tuning shifts by “increasing the discriminability in the semantic neighborhood” (Shahdloo et al., 2022). We also observed greater discriminability – after weakly constrained conceptual combination. In our study, participants were viewing only written words for concepts, so any neural change is not due to changes in sensory input (the orthography does not reflect the shape, color, etc. of the concept) but reflects a shift induced by processing words representing the concepts. What is the nature of this shift? We liken the change to that observed by Musz & Thompson-Schill (2015) who showed that EVC neural responses are more variable for words typically found in more variable contexts (e.g., ‘basket’ compared to ‘asparagus’). Their result suggests that the neural patterns underlying words change as a function of contextual variability. We propose that our semantic composition task led participants to activate a subset of features in order to complete the conceptual combination. For weakly constrained combinations, participants explored more of the semantic space, leading to less similar patterns after the composition task, akin to inducing greater contextual variability. This possibility is supported by the behavioral results, which showed that participants found it more difficult to stick to one definition for the weakly constrained combinations.

This study’s findings also add to a large body of evidence that EVC not only represents bottom-up sensory information, but is modulated by object visualization (Lee et al., 2012), visual working memory (Harrison & Tong, 2009) and retrieval of color knowledge (Hsu et al., 2012). Further, concept-level information, such as implied size (e.g., that an elephant is larger than a dog), is represented in EVC when viewing words (Borghesani, et al. 2016).

The observed EVC shift was predicted by activation levels of left ATL and right angular gyrus during composition. Specifically, participants with greater occipital pattern shifts after weakly (versus strongly) constrained combinations had greater ‘weakly > strongly constrained’ activation in left ATL and lower ‘weakly > strongly constrained’ activation (i.e., strongly > weakly) in right angular gyrus during the intermediary composition process. This fits with prior work showing modulation of top-down influences on occipital cortex by left ATL (Coutanche & Thompson-Schill, 2015) and right angular gyrus (Coutanche & Thompson-Schill, 2019). Both the ATL and angular gyrus are key hubs in the semantic memory system (Lambon Ralph et al., 2017). Here, their engagement during semantic composition predicted the extent of the EVC pattern shift for concepts underlying weakly constrained compositions. The composition process involves accessing relevant concepts, extracting features, assessing how they might be recombined or related to form a new concept, and then selecting from among several competitors (where relevant). Future studies combining fMRI and EEG (which would bring greater temporal resolution) may be able to determine which of these processes are particularly relevant to the effects we observed.

One question concerns whether EVC differences might reflect general effort or attentional processing. Typically, a general effort or attentional effect is associated with greater univariate activation (Stokes, 2011). In contrast, our univariate analyses did not reveal a difference in EVC BOLD activation during conceptual combination, consistent with the observed EVC pattern effect not reflecting greater attention. Further, the relationship established between activity in two semantic hubs (ATL and ANG; Lambon-Ralph et al., 2017) during conceptual combination and the magnitude of the EVC pattern shift is consistent with semantic processing playing a role in the effect.

### Processing of semantic compositions

Right (but not left) IFG was more active during strongly constrained than weakly constrained combinations. This was unexpected because of IFG’s role in response inhibition (Aron et al., 2003; Hampshire et al., 2009), which is expected to be greater during weakly constrained combinations (inhibiting alternative definitions of a combination; e.g., *canary bus* as a bus with small wings). We raise two considerations for this result. First, we note this effect was present in right rather than left IFG. While the left IFG has been linked with response inhibition, the right IFG is more often associated with target selection (e.g., Hampshire et al., 2009). The strongly constrained combinations may have allowed participants to hold the targeted concepts more readily via right IFG target selection. Second, left IFG activation is not always observed for multi-word phrases. For instance, Graessner et al. (2021) failed to observe IFG responses for anomalous relative to meaningful phrases and noted that the classical inhibitory N400 effect is “a lot weaker in the hemodynamic modality” (Graessner et al., 2021). We had hypothesized that the left IFG would be engaged due to its role in semantic selection and control (Solomon & Thompson-Schill, 2017; Thompson-Schill et al. 1997), and in the retrieval of weak associations (Badre & Wagner, 2007). One study found greater left IFG activation when processing unfamiliar (e.g., *house lake*) relative to familiar (e.g., *lake house*) combinations (Graves, et al. 2010). Our combinations were all novel (i.e., requiring combination for the first time), raising the possibility that a familiar versus unfamiliar contrast might be more likely to generate left IFG involvement.

Both *a priori* and exploratory results indicated that the left ANG responded more to strongly constrained than to weakly constrained combinations. This is consistent with a previous report of greater ANG activation for more meaningful word pairs (e.g., *plaid jacket* vs. *moss pony*; Price, et al. 2015), as a meaningfulness dimension is also likely reflected within our strongly constrained vs. weakly constrained contrast. Similarly, stimulating the left ANG modulates meaningful, but not meaningless, adjective-noun pairs (Price, et al. 2016). We did not find an anticipated greater ANG response to relational over attributive combinations (Coutanche, et al. 2019; Boylan et al. 2017). A possible explanation is that ANG may be driven primarily by plausibility (Graves et al. 2010; Molinaro et al. 2015; Price et al. 2015), which was similar between attributive and relational contributions. Alternatively, ANG activation may reflect its operation within the default mode network (Humphreys et al., 2015; Lambon Ralph et al., 2016; though see Graessner et al., 2021). Under this account, relational and attributive combinations are similarly difficult (leading to similar levels of ANG activation) and the easier strongly constrained combinations are accompanied by less default mode network deactivation (i.e., a stronger response).

Finally, we did not find greater ATL activation for attributive compared to relational combinations but did find differences based on constraint strength. The lack of a difference between attributional and relational combinations is consistent with a prior study that found similar magnitudes for both types of combination (Boylan, et al. 2017). Unlike Boylan et al. (2017), however, we did not find an earlier ATL BOLD response for attributive compared to relational combinations. Because the ATL is prone to signal drop-out (Devlin, et al. 2000; Ojemann, et al. 1997), a temporal signal-to-noise (tSNR) analysis was run to determine whether the null findings were due to poor signal in the ROI. Reasonable and consistent tSNR was found in both hemispheres (left ATL: range = 153 – 298, *M* = 239, *SE* = 8, 3 subjects below 200; right ATL: range = 125 – 287, *M* = 225, *SE* = 9, 7 subjects below 200), suggesting that our null findings do not reflect signal drop-out. Nonetheless, it is possible that such differences do lie within areas with impoverished signal. It is noteworthy that a recent investigation argued the ATL responds at the more general phrase level during an explicit task but not at the level of more specific ‘meaningful compositions’ (Graessner et al., 2021), suggesting that differences at this level (such as relational vs. attributional) will not be reflected in the ATL response. The exploratory wholebrain analysis identified a lateral subregion of the ATL that was more active when combining strongly constrained, compared to weakly constrained, combinations. This subregion is ventral and lateral to our ATL ROI, suggesting a ventrolateral area that responds to strongly constrained combinations, and a more medial area that is associated with EVC pattern shifts after weakly constrained combinations. This distinction is consistent with observations that the ATL is large and highly heterogeneous, with differences among its subregions (Hung et al., 2020; Skipper et al., 2011; Visser & Lambon Ralph, 2011). Future work may wish to map constraint strength in a more graded way to better understand the ATL’s organization in this respect.

The exploratory analysis also identified a cluster in the posterior cingulate cortex (pCC) as showing greater activation for strongly constrained than for weakly constrained combinations.

This region is thought to be a central hub of the default mode network and tends to deactivate during complex tasks (Leech, Braga, & Sharp, 2012; Krieger-Redwood, et al. 2016). This is consistent with weakly constrained stimuli being more complex to combine. Interestingly, recent evidence has also pointed to pCC’s involvement in creative divergent processing through increased functional coupling with prefrontal regions (Krieger-Redwood, et al. 2016).

Finally, the left parahippocampal gyrus was more active for relational than for attributive combinations. The left parahippocampal gyrus is part of a largely left-lateralized network of brain regions responsible for the storage and retrieval of semantic information (Binder, et al. 2009). It is also involved in contextual associative processing (Aminoff, Kveraga, & Bar, 2013). This finding is consistent with suggestions that relational combinations might be particularly reliant on a process of generating contexts in which the individual constituents are combined (Kenett & Thompson-Schill, 2017).

A limitation of this study is the relatively small sample size (determined based on a prior study) for the subtle differences between conceptual combination types, which can have high between-subject variability. Future studies should consider collecting additional data to account for this variability. Another limitation concerns our finding of a shift in patterns when comparing weakly and strongly constrained combinations, but not between relational and attributive combinations. This pattern of results might reflect a relational/attributive distinction that is too fine-grained to detect or a qualitative difference in how constraint-strength and sub-types of combinations are differentially represented. In either case, caution is warranted before drawing conclusions beyond the type of combination present in each finding.

To summarize, engaging in semantic composition shifted multivariate patterns associated with weakly constrained concepts in visual cortex immediately after their combination. The magnitude of this shift was predicted by activation in the left medial ATL and right ANG. Activation was greater when combining strongly constrained concepts in the lateral ATL, ANG and pCC. The left parahippocampal gyrus was more active when combining relational than combining attributive compounds, indicating differential involvement of semantic memory regions based on the type of conceptual combination being performed.

## Acknowledgments

The authors thank Drs. Natasha Tokowicz, Scott Fraundorf, and Heather D. Lucas for feedback on previous drafts. Amy Padia assisted with stimuli and data collection. Daniel Volpone, Zoe Cribbs, Xin Qian, and Paz Quinones also assisted with stimuli collection. Marc Coutanche received funding from National Science Foundation award 1947685.

